# Mammalian Neuraminidase-1 is a Critical Target of Thromboxane Induced T-Cell Immunosuppression

**DOI:** 10.64898/2025.12.01.691391

**Authors:** William W. Harless, Yunfan Li, Myron Szewczuk

## Abstract

Aspirin or celecoxib in combination with chemotherapy significantly reduced the risk for disease recurrence and improved survival after colorectal cancer surgery in the approximately 20% of patients harbouring activating mutations in the *PIK3CA* gene^1-3^. The biological mechanisms behind the anti-cancer effects of these medications are an area of intense research interest. Recently, Yang et al.^4^ found that platelet-derived thromboxane A_2_ (TXA_2_) inhibits T-cell activity through the expression of the guanine exchange factor ARHGEF1, and that aspirin was able to inhibit ARHGEF1 expression and preserve T-cell functionality and cancer immunity by blocking TXA_2_ production^4^. TXA_2_ production is inhibited by the effect of aspirin on cyclooxygenase-1 (COX-1). Celecoxib, a selective COX-2 inhibitor, does not inhibit platelet TXA_2_ production but has shown almost identical efficacy to aspirin in controlled clinical trials^3^. Here, we show that the enzyme mammalian neuraminidase 1 (NEU1) is involved in the activation and downstream signaling of the TXA_2_ receptor (TP) on T cells in response to TXA_2_. Both aspirin and celecoxib significantly inhibited TXA_2_-induced NEU1 activity and reduced ARHGEF1 expression. This regulation of the TP receptor by NEU1 may explain why aspirin and celecoxib demonstrated comparable clinical efficacy despite only aspirin being able to inhibit TXA_2_ synthesis. Targeting NEU1 may provide a novel therapeutic strategy against cancer cell metastasis by preserving a functional T-cell response against cancer.

## Introduction

After potentially curative surgical resection of a primary cancerous tumor, patients often receive chemotherapy to eradicate any occult micro-metastatic disease that may be present. The absolute curative benefit of treatment with chemotherapy after primary tumor removal for lung, breast, and colorectal cancer is approximately 5% to 15%^5^. Surgical removal of the primary tumor has been shown to increase the number of circulating tumor cells and can trigger an inflammatory response that can impair the immune response and facilitate metastasis^6^. Recent clinical studies have shown that treatment with aspirin and celecoxib added to standard chemotherapy after surgical resection of *PIK3CA* mutated colorectal cancer can significantly reduce risk for disease recurrence and improve survival^1,3^. This has prompted further studies into the biological mechanism that could explain these impressive clinical results.

An exciting research discovery that has illuminated a possible anti-cancer mechanism of aspirin found that platelet derived TXA_2_ suppressed T-cell activity by increasing the expression of the guanine exchange factor ARHGEF1^4^. ARHGEF1 was found to limit T cell effector function and disrupt the ability of T cells to control metastatic colonization in mice^4^. The report revealed that aspirin exerted its anti-cancer effects by inhibition of cyclo-oxygenase-1 (COX-1) and the production of TXA_2_ and T-cell ARHGEF1, preserving a healthy T cell response. Celecoxib inhibits cyclo-oxygenase 2 (COX-2) and has no effect on the production of thromboxane yet had almost identical efficacy to aspirin in preventing metastasis in the clinical trials cited above.

The TP receptor is a G protein coupled receptor (GPCR) that can transactivate receptor tyrosine kinases (RTK) like EGFR^7^ or the insulin receptor^8^ to induce specific cellular processes, such as cell proliferation and differentiation. A molecular organizational G protein-coupled receptor (GPCR)-signaling platform was previously discovered by us for the EGFR receptor that may be relevant to the control of the G-protein-coupled TXA_2_ receptor^9^ (Figure 1). In this molecular arrangement NEU1 and MMP9 form a complex tethered at the ectodomain of the receptor tyrosine kinase (RTK) cell surface receptor. RTK agonists induce a conformational change in the receptor, resulting in the trans-activation of the neuromedin-B receptor (NMBR), a G protein–coupled receptor (GPCR). The NMBR GPCR can also be activated directly by agonists to other GPCRs through a process of heterodimerization^10^. Here, we hypothesized that thromboxane agonists like TXA_2_ and U46619 binding to the TXA_2_ GPCR (TP) induce sialidase activity and downstream signaling depicted in Figure 1. The activated GPCR initiates Gα_i_-protein signaling which triggers the activation of MMP9 to subsequently induce the expression of NEU1. Activated MMP9 is proposed to remove the elastin-binding protein (EBP) as part of the molecular multi-enzymatic complex that contains NEU1 and protective protein cathepsin A (PPCA)^9^. Activated NEU1 hydrolyzes the α-2,3-sialyl residues linked to β-galactosides of the RTK. This desialylation process by NEU1 is predicted to remove steric hindrance of the RTK to facilitate receptor association, subsequent activation and downstream signaling. We have previously found that both aspirin and celecoxib can inhibit NEU1 expression and EGFR signaling^11^. Building on our previous discoveries and the recent discovery of the role of TXA_2_ in T-cell regulation, we investigated whether NEU1 regulates the TP G protein-coupled receptor and the subsequent expression of ARHGEF1 in T cells.

**Figure 1.**
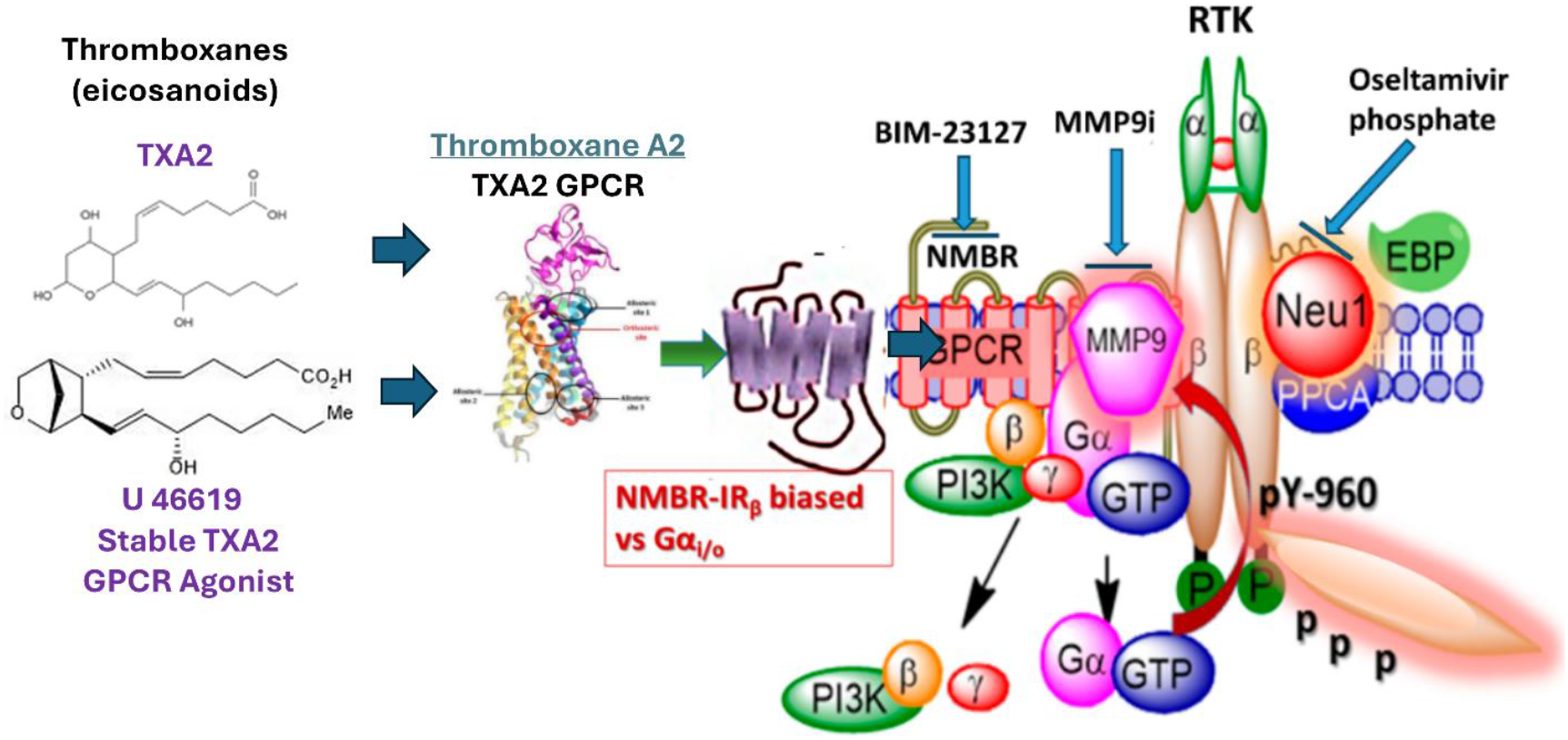
Thromboxane A2 (TXA_2_) and U46619 (stable TXA_2_ GPCR agonist) exist in a multimeric receptor complex with NMBR, RTK, and NEU1 in naïve (unstimulated) and stimulated immune cells. Here, a molecular link regulating the interaction and signaling mechanism(s) between these molecules on the cell surface uncover a biased GPCR agonist-induced RTK trans-activation signaling axis, mediated by NEU1 sialidase and the modification of RTK receptor glycosylation. The biased G-protein-coupled receptor (GPCR)-signaling platform potentiates neuraminidase-1 (NEU1) and matrix metalloproteinase-9 (MMP-9) crosstalk on the cell surface that is essential for the activation of the RTK tyrosine kinases. Notes: Activated MMP-9 is proposed to remove the elastin-binding protein (EBP) as part of the molecular multi-enzymatic complex that contains β-galactosidase/NEU1 and protective protein cathepsin A (PPCA). Activated NEU1 hydrolyzes α-2,3 sialyl residues of RTK at the ectodomain to remove steric hindrance to facilitate RTK subunits association and tyrosine kinase activation. Abbreviations: NMBR, neuromedin-B receptor; phosphatidylinositol 3-kinase; GTP: guanine triphosphate; p: phosphorylation. Citation: Taken in part from Cellular Signaling, Volume 26, Issue 6, June 2014, Pages 1355–1368. © 2014 Alghamdi et al., published by Elsevier Inc., Open access under CC BY-NC-ND license. This is an Open Access article which permits unrestricted non-commercial use, provided the original work be properly cited.

## Results

### TXA_2_, TXB_1_, and U46619 induce NEU1 sialidase activation in Jurkat Leukemia T cell line and Primary Human CD8+ Cytotoxic Killer T cells

Jurkat T cells are a well-established human T-cell leukemia cell line that possess inherent anti-tumor properties that are useful in studying how T cells can recognize and attack cancer cells, and to specifically target tumor sites^12^. Jurkat T cells have a multifaceted targeting approach to kill cancer cells, involving direct cytotoxicity, secretion of cytokines and chemokines, antibody-dependent T-cell-mediated cytotoxicity, and antibody-dependent T-cell-mediated phagocytosis^13^. Jurkat T cells can be activated via specific antigens such as proteins or peptides on the surfaces of cancer cells^14^. Interestingly, Jurkat T cells have been shown to express various growth factor receptors, including the insulin-like growth factor-I receptor (IGF-IR)^15,16^. Notably, several signaling molecules have been identified to play a critical role in the regulation of insulin-induced IR activation. Among these, the G protein-coupled receptors (GPCR) have been implicated in intracellular crosstalk pathways with IR ^8,17,18^. Since ligand binding to its GPCR can transactivate EGFR ^7^ or the insulin receptor (IGF-IR) ^9^, we investigated whether TXA_2_ binding to its GPCR, the thromboxane–prostanoid receptor (TP), can induce the activation of NEU1 sialidase in the Jurkat T-cell line, which expresses the IGF-IR.

The data depicted in Figure 2 below show that NEU1 activity is activated by exposure to U46619, **a** stable synthetic analog of TXA_2_, dose-dependently with an EC50 of 2.96 nM (**Figure 2 A-D**). We also found that specific inhibitors of MMP9, NEU1, and the neuromedin GPCR (NMBR GPCR) depicted in Figure 1, all reduced NEU1 activation in response to U46619 stimulation (**Figure 2 D**). We also tested this signaling paradigm as depicted in Figure 1 using TP agonists TXA_2_ (Figure 2 E-G). We also tested TXB_1_, which is a stable, biologically inactive metabolite of thromboxane A1 (TXA1) (Figure 2 H-J). These data support the molecular heterodimerization of the TP GPCR with the NMBR GPCR to activate MMP9 and NEU1 sialidase activity to transactivate growth factor receptors like IGF-IR expressed on Jurkat T cells. These results confirm that NEU1 is activated in this cell line after exposure to U46619, TXA2 or TXB_1_, and that the inhibition of these agonists’ induction of NEU1 is specifically inhibited by the NEU-1 inhibitor oseltamivir phosphate (OP), MMP inhibitor MMP-9i and the NMBR inhibitor BIM-23127.

**Figure 2.**
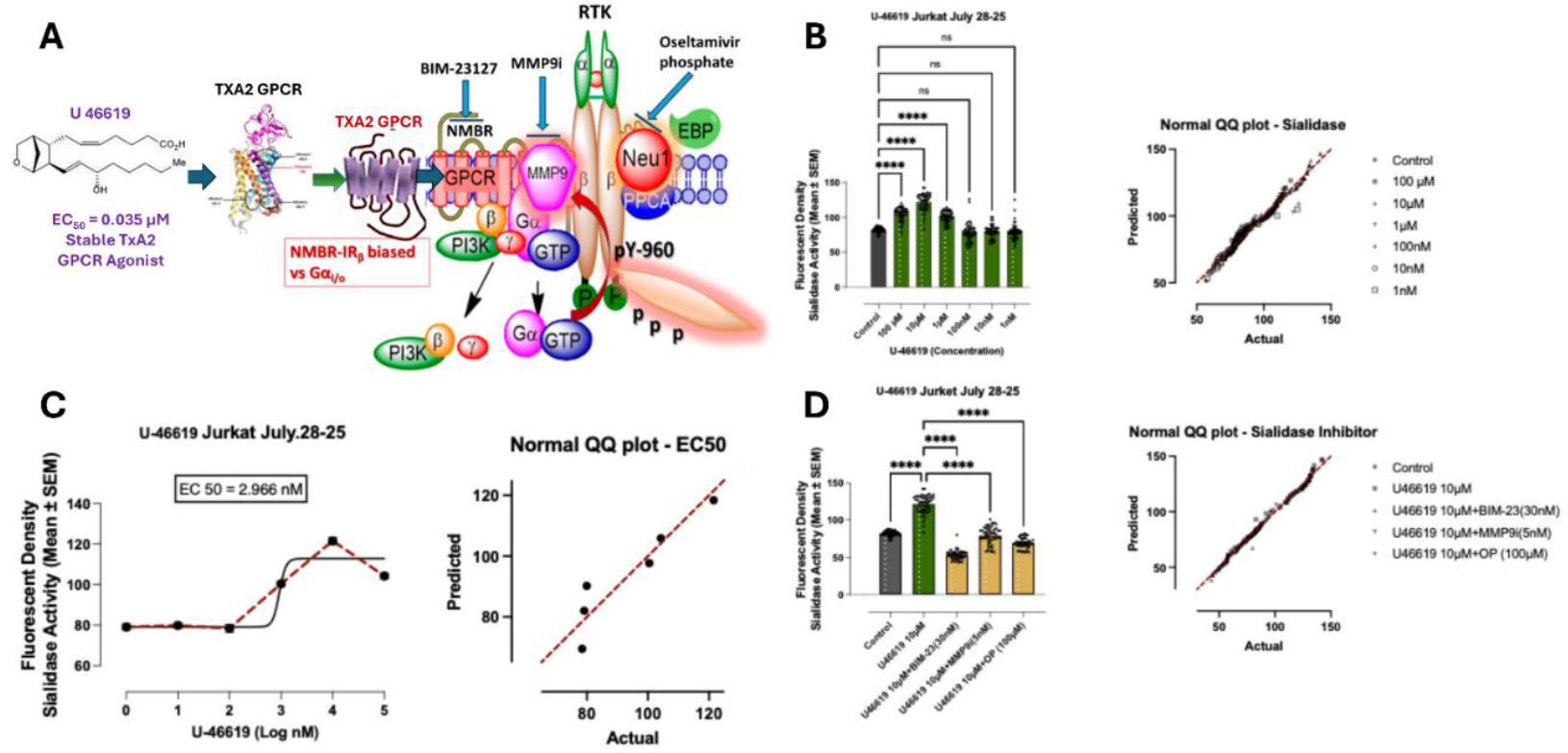
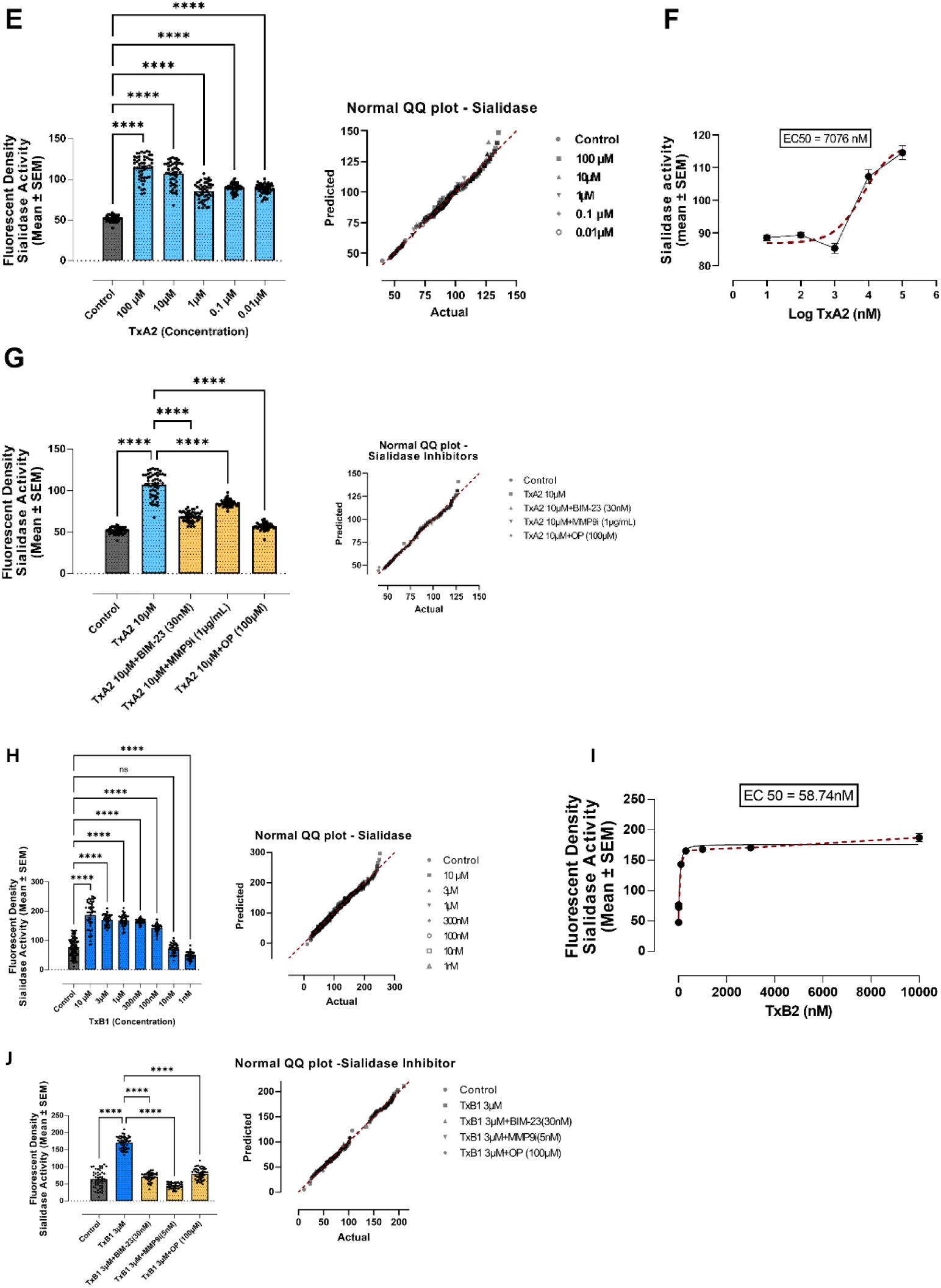
TP agonists U46619 (A-D) and TXA_2_ (E-G), as well as the metabolite of TXA_1_, TXB_1_ (H-J) increase NEU1 sialidase activity in Jurkat T cells dose-dependently, while OP, MMP9i, and BIM23 inhibitors of signaling paradigm (A) reverse TP agonists NEU1 activity. Jurkat T cells were treated with indicated agonist doses and further treated in combination with the inhibitors for one minute. 4-MUNANA substrate was added to live cells to measure sialidase activity (emission 450 nm, excitation 365 nm). The mean ± SEM (biological replicates: 5, technical replicates: 50) fluorescence density of sialidase activity demonstrates a highly significant (p < 0.0001) downregulation of sialidase activity with all three inhibitors. A normal QQ plot depicts the normality of each of the datasets, modeled by a normal distribution. Abbreviations: OP: oseltamivir phosphate; MMP9i: MMP9 inhibitor; BIM23: BIM-23127. **** p < 0.0001.

Long et al. ^19^ examined the co-expression of receptor tyrosine kinases (RTKs) and CD8+ cytotoxic T cells (CD8Ts) using a weighted gene co-expression network analysis. They found that the RTK^low^CD8T^high^ across 33 patient cancer types had the best prognosis and immune-activated microenvironment. To this end, we obtained a human primary CD8+ cytotoxic killer T cells and treated them with TP agonists U46619 and TXA2, as well as with TXB_1_. The data depicted in Figure 3 indicate a significant increase in NEU1 sialidase activity in human primary CD8+ cytotoxic killer T cells with the TP agonists TXA_2_ and U46619 but not TXB_1_ at indicated dosage. Similar to the results in the Jurkat T-cell line, OP, MMP9i, and BIM23 inhibitors reversed TP agonist TXA_2_ and U46619 induced NEU1 activity.

**Figure 3:**
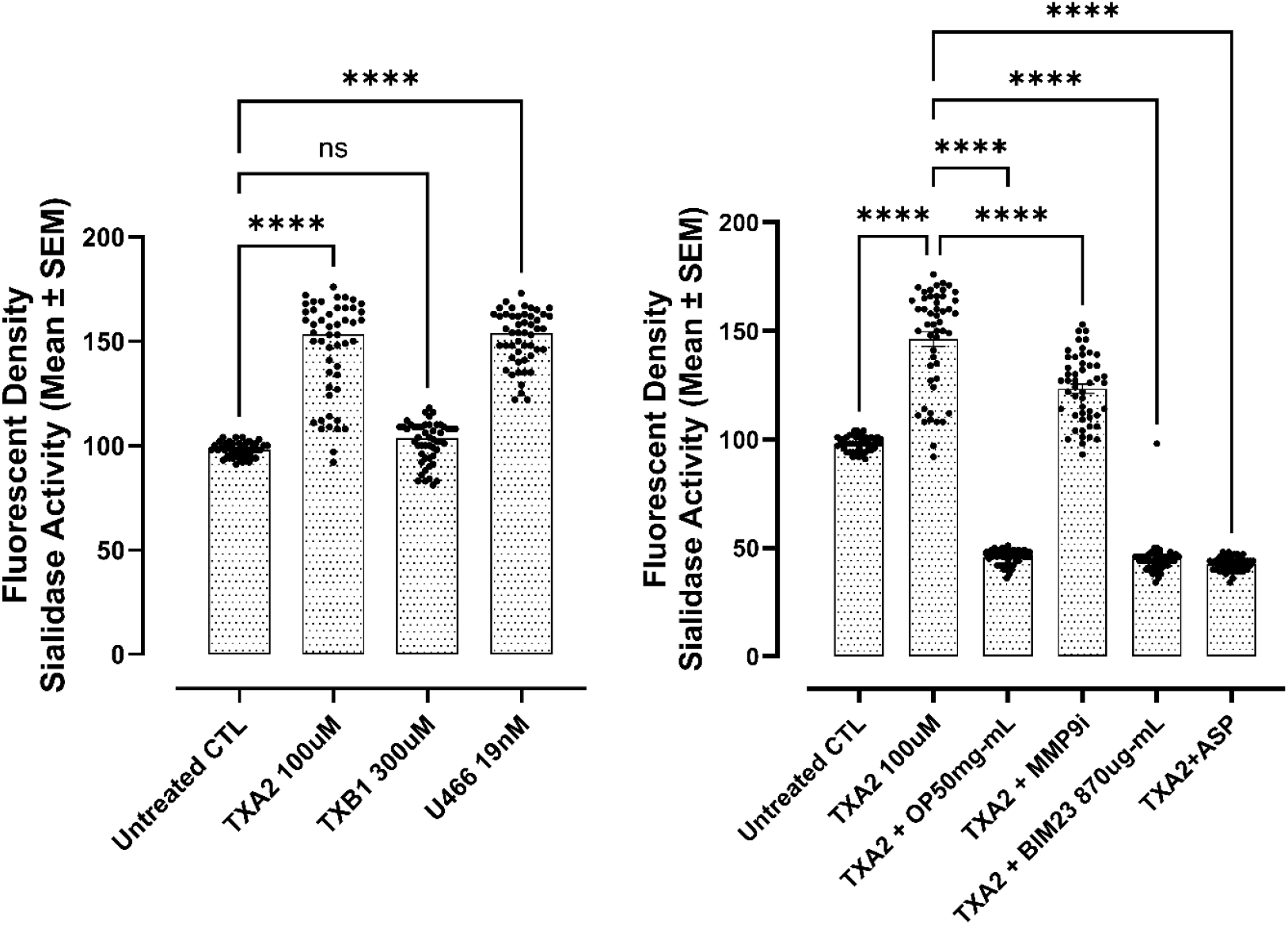
TP agonists U46619 and TXA_2_ increase NEU1 sialidase activity in human primary CD8+ cytotoxic killer T cells at indicated dosages. TXB_1_, in contrast to findings in the Jurkat T-cell line had no effect on NEU1 sialidase activity. OP, MMP9i, and BIM23 inhibitors of signaling paradigm reverse TP agonists induced NEU1 activity. Human primary cytotoxic T cells were treated with indicated agonist dosages and further treated in combination with the inhibitors for one minute. 4-MUNANA substrate was added to live cells to measure sialidase activity (emission 450 nm, excitation 365 nm). The mean ± SEM (biological replicates: 5, technical replicates: 50) fluorescence density of sialidase activity demonstrates a highly significant (p < 0.0001) downregulation of sialidase activity with all three inhibitors. Abbreviations: OP: oseltamivir phosphate; MMP9i: MMP9 inhibitor; BIM23: BIM-23127. **** p < 0.0001.

### TP GPCR Colocalizes with NEU1 on the Cell Surface of Naïve Unstimulated Jurkat T cells

The data depicted in Figure 2 demonstrated that TP receptor agonists U46619 (A-D) and TXA2 (E-G) binding to their respective GPCRs induced NEU1 sialidase activity. Here, we investigated whether the TP GPCR forms a complex with the signaling platform of NEU1 and MMP-9 crosstalk in alliance with the NMBR GPCR on the cell surface Jurkat T cells. As depicted in Figure 4, the cells were treated with equal amounts of Alexa Fluor™ 594-conjugated monoclonal antibody for the TXA_2_-GPCR (TP) and the Alexa Fluor™ 488-conjugated monoclonal anti-NEU1 antibody. The colocalization was quantified using a Pearson correlation coefficient (R-value; 0.5 < r < 0.59 = moderate positive correlation; 0.6 < r < 0.89 = strong positive correlation; r > 0.9 = near perfect correlation).

**Figure 4.**
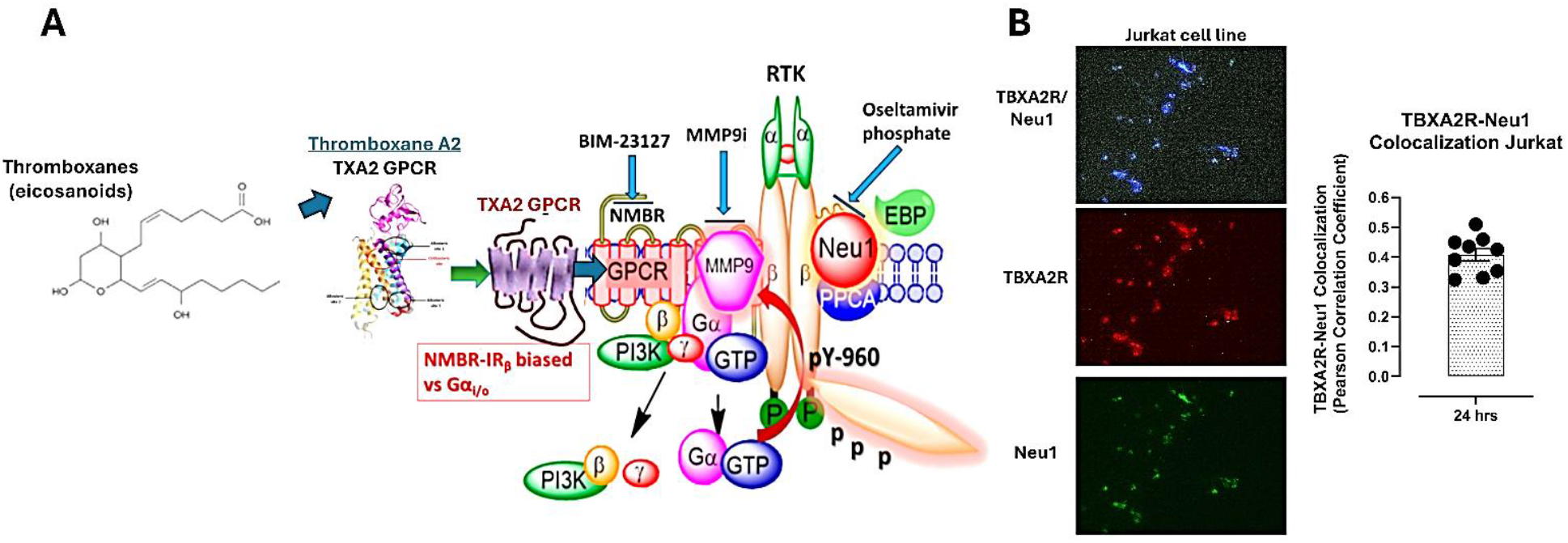
(A) TXA_2_-GPCR (TP) colocalizes with NEU1 in Jurkat T cells. (B) Cells were fixed, permeabilized, blocked, and immunostained with the specific antibodies. Cells were treated with Alexa Fluor™ 594-conjugated antibody for the TP GPCR and Alexa Fluor™ 488-conjugated NEU1 monoclonal antibody. (A) Merging of fluorescent images acquired from Zeiss M2 epifluorescent microscope (20× objective magnification) shows areas of colocalization through yellow fluorescence. Pearson correlation coefficient (R-value) measured a moderate positive correlation (0.4 < r < 0.5) between the TXA_2_-GPCR (TP) and NEU1 (biological replicates: 4, technical replicates).

For naïve Jurkat T cells, the TXA_2_-GPCR (TP) and NEU1 had a moderate positive correlation (TP: r = 0.41 ± 0.06 (Figure 3). These data confirm that the TP GPCR forms a moderate complex with NEU1 expressed on the Jurkat T-cell line.

### Celecoxib, aspirin and oseltamivir phosphate inhibition of NEU1 impede TXA_2_ and U46619 induced ARHGEF1 expression in Jurkat and Primary Human CD8+ Cytotoxic T cells

Both celecoxib and aspirin have been reported to reduce significantly disease recurrence in patients with colorectal cancer harboring the *PI3KC* mutation ^1,2^. These findings reveal a novel anticancer mechanism of both drugs in patients with *PIK3CA* mutated colorectal cancer after surgery. A recent report revealed a possible important clue to the effectiveness of aspirin in these patients with their discovery that aspirin, by inhibition of COX-1 and the production of TXA_2_, will reduce the expression of the immunosuppressive protein ARHGEF1 in T cells and preserve an effective anti-cancer T cell response^3^. Celecoxib does not affect the production of TXA_2_, yet had almost identical clinical benefit as did aspirin. Qorri et al.^11^ previously reported on a novel multimodality mechanism(s) of action for aspirin and celecoxib, specifically targeting and inhibiting NEU1 activity, as depicted in Figure 1.

Here, we investigated whether NEU1 inhibitors, oseltamivir phosphate (OP), aspirin (ASP), and celecoxib (celec), would reduce the expression of the guanine exchange factor ARHGEF1 after exposure to TXA_2_. As shown in Figure 5, TXA_2_ had a marginal, non-significant induced ARHGEF1 expression, and TXA_2_ in combination with OP, aspirin and celecoxib significantly reduced ARHGEF1 expression in Jurkat T cells after exposure to TXA_2_. It is noteworthy that the TP GPCR exhibits typical characteristics of receptor homodimerization, with specific functional selectivity. As depicted in Figure 4, TXA_2_ binding to either allosteric 1, 2 or 3 sites or orthosteric site on TP GPCR is predicted to have a biased functional homodimerization with NMBR GPCR on RTK to induce downstream activation of ARHGEF1 expression. Here, the data suggest an allosteric binding of TXA_2_ to TP affects TP-homodimerization with NMBR with minimal downstream activation.

**Figure 5.**
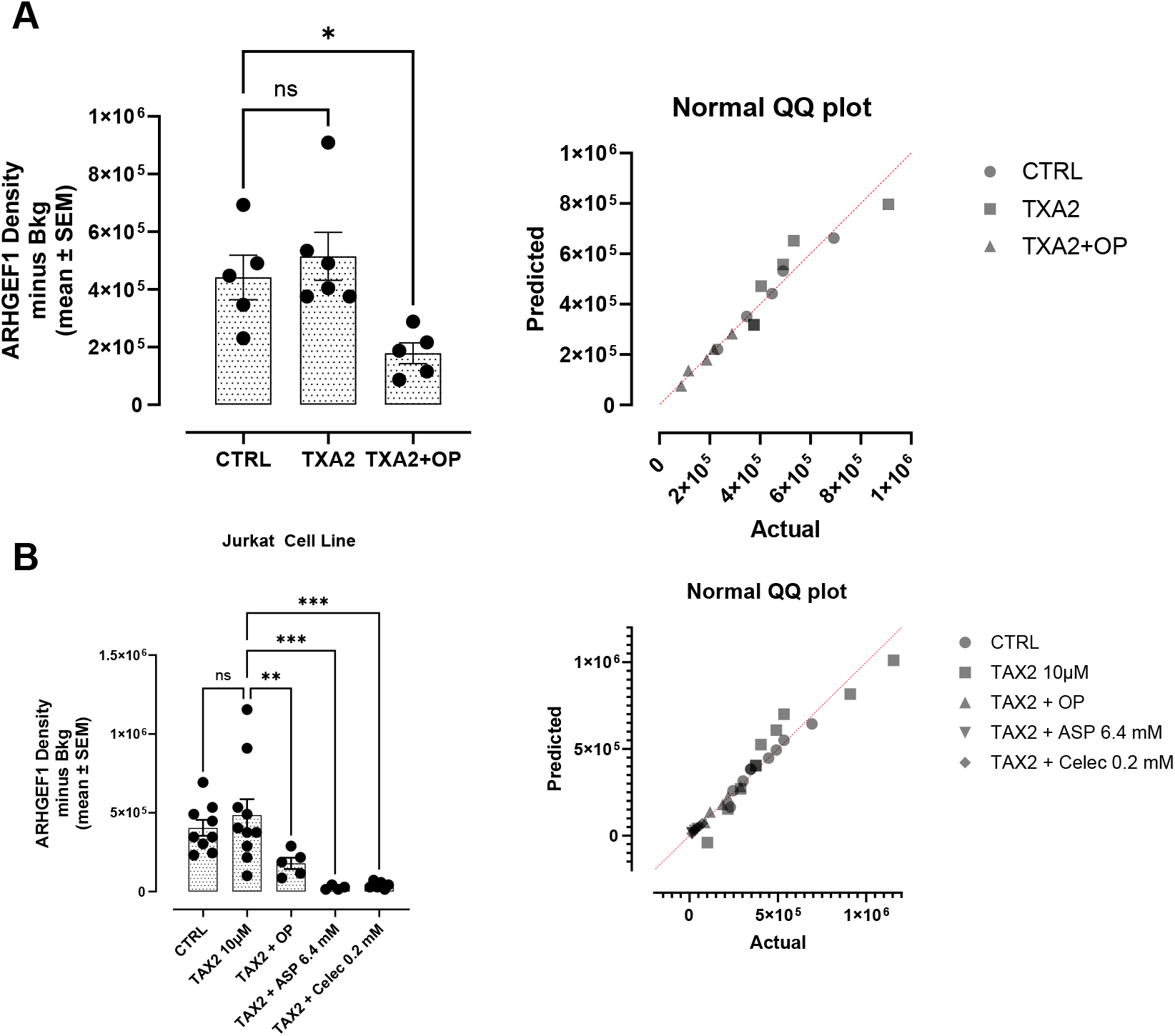
Analyses of Jurkat T cells for ARHGEF1 expression following 24-hour exposure to TXA_2_ in a combination of ASA, celecoxib and OP cocktail using immunocytochemistry. (A) Jurkat T cells were plated at 150,000 cell/mL and left untreated, treated with TXA2 (10 µM) in combination with OP (300 µg/mL), ASP (6.4 mM) or celecoxib (0.2 mM) for 24 hours. Cells were immunostained for ARHGEF1 expression using rabbit anti-ARHGEF1 and secondary goat antirabbit -Alexa 594. The results are depicted as a scatter plot of data visualization using dots to represent the indicated ARHGEF1 expression values obtained from 2 independent experiments of multiple images (n = 4–10). The mean density staining corrected for background (Bkg) ± S.E.M. for indicated ARHGEF1 expression values is indicated for each group. The density staining values of each group were compared to the untreated control as well as TXA_2_ in combination with inhibitor cocktail by ANOVA using the uncorrected Fisher’s LSD multiple comparisons test with 95% confidence with indicated asterisks for statistical significance. (B) Immunocytochemistry of untreated and TXA2 (10 μM) treated Jurkat T cells for 24 hrs in combination with ASP, celecoxib and OP. Data are presented as the percentage of cells expressing ARHGEF1corrected for background autofluorescence. ns, not significant, *p ≤ 0.01, **p ≤ 0.001, ***p ≤ 0.0001. Abbreviations: ASP, aspirin (acetylsalicylic acid); celec, celecoxib; OP, oseltamivir phosphate.

We also tested these NEU1 inhibitors on U46619-induced ARHGEF1 expression in primary human CD8+ cytotoxic killer T cells. The data depicted in Figure 6 show that U46619 had a significant increase in ARHGEF1 expression (p ≤ 0.01), and U46619 in combination with OP, aspirin and celecoxib significantly reduced ARHGEF1 expression (p ≤ 0.0001). Interestingly, ARHGEF1 expression is predominantly expressed by specific populations of tumor-infiltrating myeloid cells and regulatory myeloid cells, where it plays a pro-tumorigenic and T cell suppressive role^20^. ARHGEF1-expressing myeloid cells suppress T cell function by metabolizing the local L-arginine, which is essential for T cell activation and proliferation^21^.

**Figure 6.**
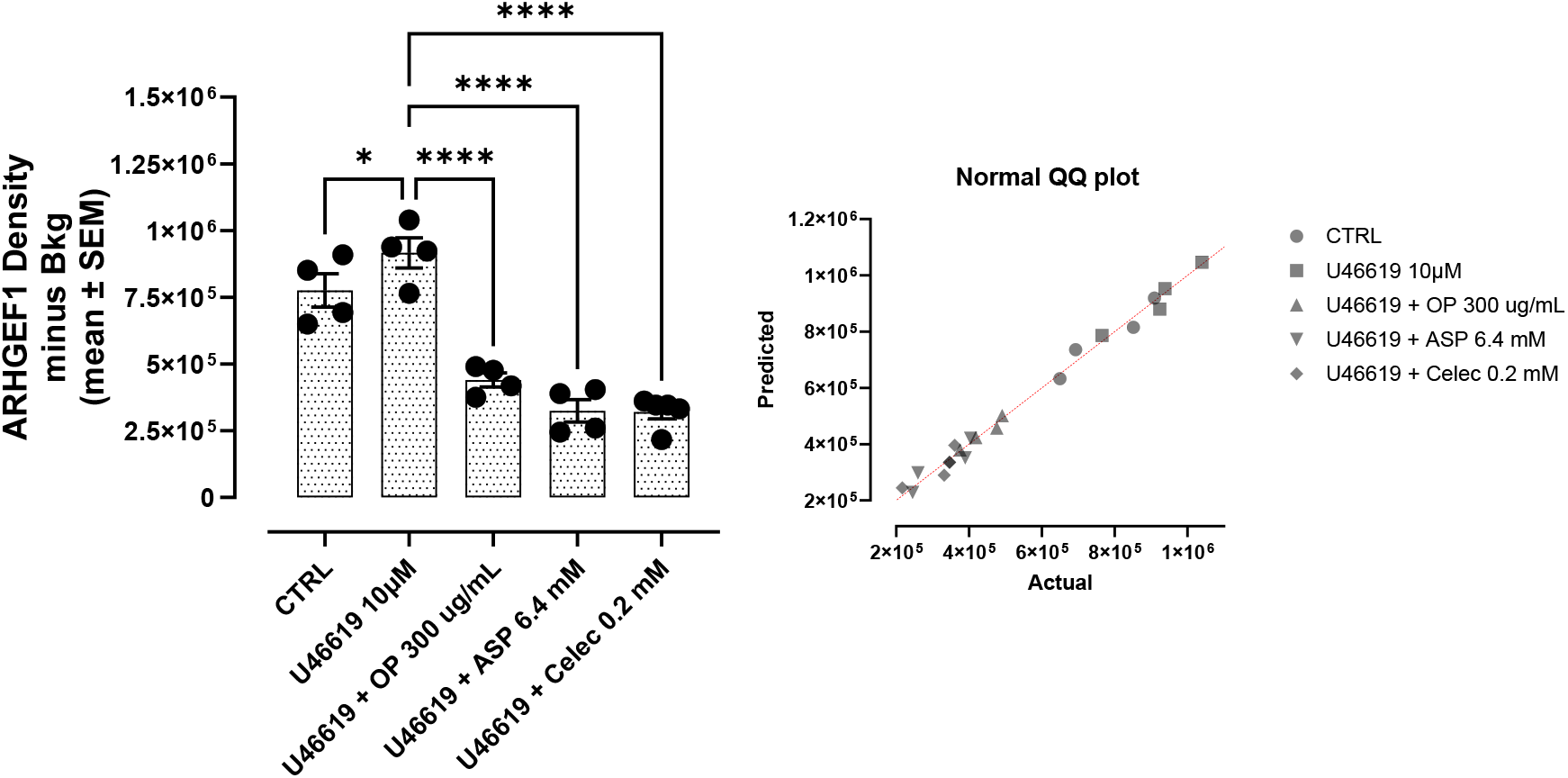
Analyses of human primary CD8+ cytotoxic killer T cells for ARHGEF1 expression following 4-hour exposure to U46619 in a combination of ASA, celecoxib and OP cocktail using immunocytochemistry. (A) Human primary CD8+ cytotoxic killer T cells were plated at 150,000 cell/mL and left untreated, treated with U46619 (10 µM) in combination with OP (300 µg/mL), ASP (6.4 mM) or celecoxib (0.2 mM) for 4 hours. Cells were immunostained for ARHGEF1 expression using rabbit anti-ARHGEF1 and secondary goat anti-rabbit -Alexa 594. The results are depicted as a scatter plot of data visualization using dots to represent the indicated ARHGEF1 expression values obtained from 2 independent experiments of multiple images (n = 4–10). The mean density staining corrected for background (Bkg) ± S.E.M. for indicated ARHGEF1 expression values is indicated for each group. A normal QQ plot depicts the normality of each of the datasets, modeled by a normal distribution. The density staining values of each group were compared to the untreated control as well as U46619 in combination with inhibitor cocktail by ANOVA using the uncorrected Fisher’s LSD multiple comparisons test with 95% confidence with indicated asterisks for statistical significance. Data is presented as the percentage of cells expressing ARHGEF1corrected for background autofluorescence. ns, not significant, *p ≤ 0.01, ***p ≤ 0.0001. Abbreviations: ASP, aspirin (acetylsalicylic acid); celec, celecoxib; OP, oseltamivir phosphate.

## Discussion

Platelet derived TXA_2_ triggers T cell immunosuppression dependent on the synthesis of the guanine exchange factor ARHGEF1^4^, suppressing T cell receptor-driven kinase signalling, proliferation and effector functions. The report has shown that T cell-specific conditional deletion of the *Arhgef1* gene in mice increased T cell activation at the metastatic site, thereby provoking an immune-mediated rejection of lung and liver metastases^4^. TXA2 is produced by platelets under the control of cyclooxygenase 1 (COX-1) but not cyclooxygenase-2 (COX-2). Aspirin blocks both COX-1 and COX-2, but celecoxib specifically blocks COX-2 and does not affect TXA_2_ production. Both celecoxib and aspirin reduced significantly disease recurrence in patients with colorectal cancer harboring the *PIK3CA* mutation^1-3^. These benefits were surprising and staggering: Low-dose aspirin for three years after surgical removal of the primary tumor reduced the risk of colorectal cancer recurrence by around 50% in patients with *PIK3CA* mutations (hazard ratio ∼0.49)^2^. Similarly, patients with *PIK3CA* mutated colorectal tumors receiving celecoxib after surgery had their risk of dying reduced by about 50% to 60%^3^.

Here we show that the TP GPCR on the human Jurkat T cell line and CD8 + cytotoxic T cell line is regulated by mammalian neuraminidase-1 (NEU1). Blocking NEU1 with either aspirin, celecoxib or oseltamivir phosphate reduced significantly the expression of ARHGEF1. This reduction in the expression of ARHGEF1 would be predicted to be important in the preservation of a T cell mediated immune response as previously noted. The regulation of the T cell TP receptor by NEU1 may solve the riddle of why both aspirin and celecoxib had virtually identical clinical results despite only aspirin being able to inhibit TXA2 expression.

A potentially conflicting finding with our results on the critical role of NEU1 regulating the TP receptor and ARHGEF1 expression is that administering the TXA_2_ analog U46619 overcame aspirin’s effectiveness in preventing metastasis in mice^4,22^. This observation suggests that it is the inhibition of COX-1 and the production of TXA_2_ that is a primary anti-cancer mechanism of action for aspirin. But this experimental result does not necessarily contradict our finding of the central role played by NEU1 in the regulation of the TP receptor and ARHGEF 1 expression. U46619 is a chemically stable analogue of TXA_2_ that produces sustained TP receptor engagement in distinct contrast to the highly labile and spatially restricted signaling of endogenous TXA_2_ in vivo^23^. A **sustained** TP receptor engagement would be predicted to overcome the effect of aspirin on the suppression of TP receptor signaling by NEU1. In our experiments we were able to completely shut down the expression of ARHGEF1 using NEU1 inhibitors after stimulation with TXA_2_ but only partially able to do so after stimulation of the cells using U46619.

TXA_2_ is significantly upregulated after cancer surgery, particularly in GI malignancies, and the use of aspirin to target this upregulation after surgical removal of a primary tumor has previously been proposed^24^. Upregulated TXA_2/_TP receptor signaling would be predicted to be host immunosuppressive, particularly right after surgery and the time when circulating tumor cells might be shed and the risk for metastatic spread of these cells is enhanced due to immune suppression after surgery^6^.

*PIK3CA* mutant cancers have increased levels of expression of COX-2 and upregulated expression of prostaglandin E2 (PGE_2)_ ^25^. PGE_2_ signalling inhibits cytotoxic T lymphocyte (CTL) and natural killer (NK) cell activity, changes macrophage polarization toward a pro-tumour M2 phenotype, and impairs dendritic cell maturation and antigen presentation fostering immune suppression ^26^. *PIK3CA*-mutated tumors recruit myeloid-derived suppressor cells (MDSCs) that suppress the activity of cytotoxic T cells via the arachidonic acid (AA) metabolism pathway^27^. PIK3CA expression in tumors was found to decrease the number of CD8^+^ T cells but to increase the number of inhibitory myeloid cells following immunotherapy ^28^. Myeloid derived suppressor T cells presence in the circulation is predictive of disease recurrence after cancer surgery^29^.

In contrast to the distinct immunosuppression triggered by *PIK3CA* mutated cancers, there is evidence that these cancers may be potentially highly susceptible to a host immune response. In a syngeneic pancreatic cancer model, genetic loss of *PIK3CA* increased tumor T cell infiltration and led to tumor regression that depended on T cells^30^. Furthermore, *PIK3CA* deficient tumors expressed higher MHC-I and costimulatory molecules, thus becoming more susceptible to immune clearance^30^. These results indicate that *PIK3CA*-mutated cancers may be characterized by the proverbial double-edged sword. The biology of these cancers upregulates immunosuppressive mechanisms in the host to facilitate their survival and metastasis, but they may also be distinctly vulnerable to an effective host immune response. In this context, *PIK3CA* mutated cancers would be predicted to be highly sensitive to therapeutic manipulation of TXA_2/_TP receptor signaling, particularly after surgical removal of the primary tumor and the time when an effective host immune response is critical to the eradication of any surviving cancer cell population.

Whether the impressive clinical results using aspirin and celecoxib after surgery in the *PIK3CA* mutated colorectal cancer patients could be extended to other *PIK3CA* mutated cancers is currently unknown. This is not a trivial matter as the results in the colorectal cancer patients using inexpensive and relatively safe oral drugs were dramatic and offered a curative treatment with minimal side effects. *PIK3CA* mutations occur at varying frequencies in different forms of cancer. Notably, these genetic alterations are present in approximately 30 to 35 percent of cases involving breast, endometrial, and cervical cancers^31^. In addition to these, *PIK3CA* mutations have also been identified in cancers such as colorectal, gastric, ovarian, and lung cancer, among others^31^. This broad distribution highlights the significance of *PIK3CA* mutants in the molecular landscape of various malignancies. The therapeutic targeting of NEU1 as an anti-cancer treatment should be further explored in clinical trials, particularly in patients harboring *PIK3CA* mutations. This treatment would be predicted to be particularly important after surgical removal of the primary tumor and the time when effective treatment can make the difference between disease eradication or metastatic spread of the cancer^32,33^.

## Methods

### Cell lines

Human Jurkat leukemia T cells (Clone E6-1, TIB-152 ™) obtained from the American Type Culture Collection, Manassas, VA 20110-2209, USA, are an immortalized human T-lymphocyte cell line derived from a patient with T cell acute lymphoblastic leukemia (T-ALL). Human CD8+ cytotoxic killer T cells are primary cells obtained from blood (Cat No. T4121, Applied Biological Materials, Inc., Richmond, BC, Canada). Cells were grown in culture media containing Dulbecco’s Modified Eagle’s Medium (DMEM) (Gibco, Rockville, MD, U.S.A.) supplemented with 10% fetal bovine serum (FBS) (HyClone, Logan, UT, U.S.A.) and 5 μg/mL plasmocin (InvivoGen, San Diego, CA, U.S.A.) as a prophylactic inhibitor of mycoplasma. Cells with single passage were maintained in an incubator with 5% CO_2_ at 37°C until 75% confluence was reached.

### Reagents

U-46619 (Item No. 16450, Formal Name, (5Z)-7-[(1R,4S,5S,6R)-6-[(1E,3S)-3-hydroxy-1-octen-1-yl]-2-oxabicyclo[2.2.1]hept-5-yl]-5-heptenoic acid) (Cayman Chemical, Ann Arbor, Michigan 48108 USA) is a stable analog of the endoperoxide prostaglandin H2, and a TP receptor agonist.1 It exhibits properties similar to thromboxane A2 (TXA2), causing platelet shape change and aggregation, and contraction of vascular smooth muscle. 15(S)-Pinane Thromboxane A2 (Item No. 19020, Formal Name, (5Z)-7-[(1S,2R,3R,5S)-3-[(1E,3S)-3-hydroxy-1-octen-1-yl]-6,6-dimethylbicyclo[3.1.1]hept-2-yl]-5-heptenoic acid) (Cayman Chemical, Ann Arbor, Michigan 48108 USA) is a stable analog of TXA2. Thromboxane B1 (Item No. 10006610, Batch No. 0644291, Formal Name 9,11,15-trihydroxy-thrombox-13-en-1-oic acid) (Cayman Chemical, Ann Arbor, Michigan 48108 USA) is a non-enzymatically derived, stable, inactive metabolite of TXA2, and positively correlates with platelet COX-2 activation. Rabbit anti-ARHGEF1 antibody (Cat # PA578821, Life Technologies, Inc.) was obtained from Thermo Fisher, Canada at a dilution of 1 μg (per 1×10^6^ cells), followed by Alexa-594 conjugated goat anti-rabbit IgG for 30 minutes at 4°C using 5-10 μg (per 1×10^6^ cells) dilution. Rabbit monoclonal anti-TBXA2 receptor (Cat # MA557283, Life Technologies, Inc.) was obtained from Thermo Fisher, Canada, at a dilution of 1:100, followed by Alexa-594 conjugated goat anti-rabbit IgG for 30 minutes at 4°C using 5-10 μg (per 1×10^6^ cells) dilution.

### Inhibitors

Oseltamivir phosphate (OP) (>99% pure OP, batch No. MBAS20014A, Solara Active Pharma Sciences Ltd., New Mangalore-575011, Karnataka, India), a broad-spectrum sialidase inhibitor that inhibits NEU1, was used at a concentration of 300 μg/mL, as previously reported by Baghaie et al. ^28^. Acetylsalicylic acid (>99% pure, Sigma-Aldrich, Steinheim, Germany) was dissolved in dimethyl sulfoxide (DMSO) to prepare a 5000 mM stock solution and stored in aliquots at −20°C. The highest used concentration of aspirin contains less than 0.5% v/v of DMSO in 1× PBS at a pH of 7. Celecoxib (Marcan Pharmaceuticals Inc., Ottawa, ON, Canada) was dissolved in DMSO to prepare a stock solution of 52.44 mM and stored in aliquots at −20°C.

### Live Cell Sialidase Assay

Jurkat T-cells at a density of 100,000 cells/well were centrifuged at 900 rpms on 12 mm circular glass slides in sterile 24-well plates. 2 μL of 0.318 mM 2'-(4-methylumbelliferyl)-α-D-N-acetylneuraminic acid sodium salt hydrate (4-MUNANA; ≥95% pure, M8639, Sigma-Aldrich) substrate in Tris-buffered saline (TBS, pH 7.4) was added to a coverslip alone as a control, with 2 μL of 30 ng/mL TXA2 (, Cedarlane), or with 2 μL of TXA2 and 2 μL of aspirin or celecoxib at the indicated concentrations. 4-MUNANA is hydrolyzed by sialidase activity on the cell surface, releasing 4-methyl-lumbelliferone (4-MU), which has a fluorescence emission at 450 nm (blue color) following excitation at 365 nm. Fluorescent images were taken at 20× after 1–2 minutes of treatment using the Zeiss M2 Imager epi-fluorescent microscope. The mean fluorescence of 50 multi-point replicates surrounding the cells was quantified using the ImageJ software.

### Co-Localization

Jurkat T-cells at a density of 100,000 cells/well were centrifuged at 900 rpms on 12 mm circular glass slides in sterile 24-well plates. Cells were fixed with 4 μg/mL paraformaldehyde (PFA) for 24 h at a 5 ?C refrigerator. Subsequently, cells were permeabilized with 0.1% TritonX in PBS (PBST) for 5 min and then blocked with 4% BSA in 0.1% Tween-TBS in the fridge for 1 h. Cells were treated with monoclonal rabbit anti-TXA2 receptor at 4°C for 24 hrs, washed 3x with PBS and followed with Alexa-488 conjugated mouse anti-NEU1 antibodies at 4°C for 24 hrs. Coverslips with stained cells were mounted on microscope slides onto 3 μL of DAPI fluorescent mounting medium. Slide images were observed using ZeissM2 epi-fluorescent microscopy (40× magnification, Carl Zeiss Canada Ltd., M3B 2S6 Toronto, Canada), capturing images under green (488 nm) and red (594 nm) channels. The Pearson correlation coefficient quantifies protein colocalization in the acquired images, and the results were expressed as a percentage determined using AxioVision software, Rel. version 4.6. Pearson’s correlation coefficient measures the linear proximity and association between two variables. A correlation value of 0.5 to 0.7 between the two variables would indicate that a significant and positive relationship exists between the two.

### Immunocytochemistry

Jurkat T cells plated at 150,000 cell/mL were centrifuged at 900 rpms on 12 mm circular glass slides in sterile 24-well plates. Cells were left untreated, treated with TXA2 (10 μM) or in combination with OP (300 μg/mL), ASP (6.4 mM) or celecoxib (0.2 mM) for 24 hours at 37°C. Cells were washed and fixed with 4% paraformaldehyde (PFA) for 30 minutes, followed by permeabilization with 0.1% TritonX in PBS (PBST) for 10 minutes. Cells were blocked with 4% BSA in PBST for 1 hour at 4°C. Following blocking, cells were washed with 1× PBS 3× for 10 minutes, followed by the addition of rabbit monoclonal anti-ARHGEF1 diluted in 4% BSA in PBST in a 1:20 ratio overnight at 4°C. Rabbit serum were used and diluted in the same ratio as primary antibodies for isotype controls. Cells were washed 3× for 5 minutes with 1× PBS and incubated for 1 hour with Goat anti-rabbit AlexaFluor Plus 594 (A32740, Invitrogen). Cells were then washed 3× for 10 minutes. Coverslips with attached cells were inverted on a droplet of mounting media containing 4′,6-diamidino-2-phenylindole (DAPI; VECTH1200, MJS BioLynx Inc., P.O. Box 1150, 300 Laurier Blvd., Brockville, ON K6V 5W1, Canada) to visualize the nuclei and sealed. The stained cells were visualized by epifluorescence microscopy at 20× mag.

### Statistical Analysis

All statistical analyses were performed using GraphPad Prism 10 software and are presented as the mean ± standard error of the mean (SEM). Comparisons between groups from two to three independent experiments were made by one-way analysis of variance (ANOVA) at 95% confidence using the uncorrected Fisher’s LSD multiple comparisons test with 95% confidence. Asterisks denote statistical significance.

#### Patents

M.R.S. reports patents for the use of NEU1 sialidase inhibitors in cancer treatment (Canadian patent no. 2858,246; United States patent no. US2015/0064282 A1; European patent no. 1874886.2; Chinese patent no. ZL201180076213.7; German patent no. 602011064575.7; Italian patent no. 502020000014650; UK patent no. 2773340; Swedish patent no. 2773340; Spanish patent no. 773340; Switzerland patent no. 2773340; and French patent no. 2773340). M.R.S. reports a patent for using oseltamivir phosphate and analogs thereof to treat cancer (International PCT patent no. PCT/CA2011/050690). W.W.H. and M.R.S. report a patent on a method to improve the effectiveness of anti-cancer therapies by exposing them to an inflammatory stimulus prior to treatment (Canadian patent no. PCT/CA2017/050765, pending). W.W.H. and M.R.S. report a patent on the compositions and methods for cancer treatment (Cana-dian patent no. PCT/CA2017/050768, pending). W.W.H. and M.R.S. report patents 55983477-9CN and CAN_DMS_150056368.1 on the COMPOSITIONS AND METHODS FOR THE TREATMENT OF CORONAVIRUS INFECTION AND RESPIRATORY COM-PROMISE.

#### Clinical Trials USFDA

W.W.H. and M.R.S. have US FDA clinical trial approval to test OP in patients with pancreatic cancer (clinical trial number #173874).

## Abbreviations

PFA: paraformaldehyde
BSA: Bovine Serum Albumin
SEM: mean ± standard error of the mean
ANOVA: one-way analysis of variance
FBS: Fetal Bovine Serum
MMP9i: metalloproteinase inhibitor
MET: mesenchymal-epithelial transition
LPS: lipopolysaccharide

## Author Contributions

YL, WWH, and M.R.S. analyzed the literature and contributed to writing and revising the manuscript. All authors made a significant contribution to the work reported, whether that is in the conception, study design, execution, acquisition of data, analysis and interpretation, or all these areas; took part in drafting, revising, or critically reviewing the article; gave final approval of the version to be published; have agreed on the journal to which the article has been submitted; and agree to be accountable for all aspects of the work. All authors have read and agreed to the published version of the manuscript.

## Informed Consent Statement

Not applicable.

## Data Availability Statement

All data needed to evaluate the paper’s conclusions are present. The preclinical data sets generated and analyzed during the current study are not publicly available but can be obtained from the corresponding author upon reasonable request. The data will be provided following the review and approval of a research proposal, Statistical Analysis Plan, and execution of a Data Sharing Agreement. The data will be accessible for 12 months for approved requests, subject to possible extensions; contact for more information on the process or to submit a request.

## Acknowledgments

Y.L. is the recipient of the 2022-2023 Dean’s Honour List, the E.D. Merkley Prize in Mathematics, and the M.C. Urquhart Book Prize in Economics. All authors have read and agreed to the published version of the manuscript. All authors acknowledge the educational and scholarly alliance of the Graduate Program in Experimental Medicine and the Health Sciences Research program.

## Peer review information

Nature thanks Reviewers (named) and the other, anonymous, reviewer(s) for their contribution to the peer review of this work.

## Reprints and permissions information

is available at http://www.nature.com/reprints.

